# Systematic Characterization of LSD metabolites in *C. elegans* by ultra-high performance liquid chromatography coupled with high-resolution tandem mass spectrometry

**DOI:** 10.1101/2023.06.19.545563

**Authors:** Christiane Martins de Vasconcellos Silveira, Vanessa Farelo dos Santos, Isis Moraes Ornelas, Beatriz de Sá Carrilho, Matheus Antônio Vieira de Castro Ventura, Henrique Marcelo Gualberto Pereira, Stevens Kastrup Rehen, Magno Junqueira

## Abstract

Psychedelic compounds have gained renewed interest for their potential therapeutic applications, but their metabolism and effects on complex biological systems remain poorly understood. Here, we present a systematic characterization of LSD metabolites in the model organism *Caenorhabditis elegans* using state-of-the-art analytical techniques. By employing ultra-high performance liquid chromatography coupled with high-resolution tandem mass spectrometry (UHPLC-HRMS/MS), we identified and quantified a range of LSD metabolites, shedding light on their metabolic pathways and offering insights into their pharmacokinetics. Our study demonstrates the suitability of *C. elegans* as a valuable model system for investigating the metabolism of psychedelic compounds and provides a foundation for further research on the therapeutic potential of LSD. These findings contribute to the growing body of knowledge in the field and highlight the importance of advanced analytical methodologies in elucidating the effects of psychedelic substances on biological systems.

## Introduction

Lysergic Acid Diethylamide (LSD), a psychedelic first synthesized on 1938, by the Swiss chemist Albert Hofmann in the Sandoz (now Novartis) laboratories in Basel, Switzerland [1], was the subject of studies of clinical and psychiatric effects [2,3,4,5,6,7]. Due to the hallucinogenic properties, it was widely used for recreational purposes, but its use was prohibited in the 1960s [8,9]. Currently, LSD has been reintroduced in research since LSD-assisted psychotherapy is studied as an alternative for treatment of psychiatric illnesses and disorders, such as anxiety and depression [2,3,7]. Depression, one of the most common psychiatric disorders, affects more than 300 million people around the world [10,11,12]. Some of the symptoms include mood swings, anhedonia, psychomotor changes, and suicidal tendencies [11]. Treatments with many antidepressants have been used in recent decades, but a low efficacy of 30% to 40% among patients, as well as a time interval of 3 to 4 weeks for the appearance of results, have stimulated the search for new options [10,12,13]. In addition to LSD, other serotonergic psychedelics, such as psilocybin, mescaline and dimethyltryptamine, have also shown positive and promising results, since, unlike traditional antidepressants, such as monoamine oxidase inhibitors and selective serotonin reuptake inhibitors, they act as full or partial serotonin 5-HT2A receptor agonists. [10,11,12,13,14].

The possibility of including LSD as a psychotherapy option requires information about its metabolism, pharmacology, and pharmacokinetics. The development of studies in humans address ethical issues such as the patient’s vulnerability to addiction, and evaluation of risks and benefits, which must be well described, supervised, and managed precisely [15,16]. Thus, the use of the nematode *Caenorhabditis elegans* (*C. elegans*), a widely applied model in neurological studies [17,18,19,20,21], and represents an alternative to corroborate studies with humans. Recent contributions have used *C. elegans* model in the study of axonal regeneration [22], autism [23], neuro-intestinal ferritin regulation [24], Alzheimer’s gene expression and neuronal apoptosis [25], and in neural plasticity [26]. Some of the advantages of working with the model are fast reproduction rate, easy maintenance in the laboratory, a small nervous system, and, mainly, the entire genome sequenced [17,18,19,20,21]. Due to the high genetic homology with humans, ranging from 60% to 80%, genetic manipulation of the nematode has been explored in several studies on aging and human diseases, both at the metabolic and genomic level [17,18,19,20,21]. In behavioral and metabolic studies with substances of abuse, there are few established models using matrix with *C. elegans*, with records referring only to alcohol, cocaine, methamphetamine and nicotine [18,27,28]. To the authors knowledge, there are metabolic studies with LSD only in rats, mice, guinea pigs, cats, and humans [5,29,30].

The aim of the study was to identify LSD metabolites and their production rate, employing *C. elegans* as a model, by performing ultra-high performance liquid chromatography coupled with high-resolution tandem mass spectrometry (UHPLC-HRMS/MS). The expected low concentrations of LSD and its derivatives require increasingly sensitive and accurate identification and quantification methods. The advances and widespread use of UHPLC-HRMS/MS with electrospray ionization (ESI) permits its application in such field, mainly with methods including a second stage of mass fragmentation [31,32].

## Materials and methods

### Chemicals and Reagents

LSD and the internal standard (ISTD) LSD-D3 were purchased from Sigma-Aldrich (São Paulo, Brazil). *C. elegans* were obtained from Caenorhabditis Genetics Center (CGC), Minnesota, USA. Ultrapure water (18.2 MΩ·cm) was obtained from a Millipore Milli-Q purification system (Billerica, MA, USA). Methanol (HPLC grade), formic acid 98-100%, and ammonium formate were acquired from Tedia (Fairfield, OH, USA), Merck (Darmstadt, Germany) and Spectrum Chemical (Gardena, CA, USA), respectively. Potassium phosphate monobasic (KH_2_PO_4_), sodium phosphate dibasic (Na_2_HPO_4_), cholesterol, magnesium sulfate (MgSO_4_) and streptomycin sulfate salt were purchased from Sigma Aldrich (São Paulo, Brazil). Calcium chloride (CaCl_2_) and sodium chloride (NaCl) were purchased from Merck (Darmstadt, Germany). Agar bacteriological, bacto peptone and potassium phosphate dibasic were purchased from Thermo Scientific (Massachusetts, USA).

### Sample preparation

The preparation of the samples for analysis followed three steps, as represented in figure 1: (a) worms’ growth, (b) extract preparation with LSD, and (c) addition of the ISTD LSD-D3 and analysis by UHPLC-HRMS/MS. **Worms growth and treatment:** Experiments were performed with wild type strain, N2 Bristol. Animals were cultivated according to well stablished protocols (Brenner, 1974). For maintenance, animals were grown at 15°C or 20°C, on NGM (nematode growth media) plates seeded with *Escherichia coli* (OP50-1) and supplemented with 100 μg/L streptomycin.

**Figure 1.**
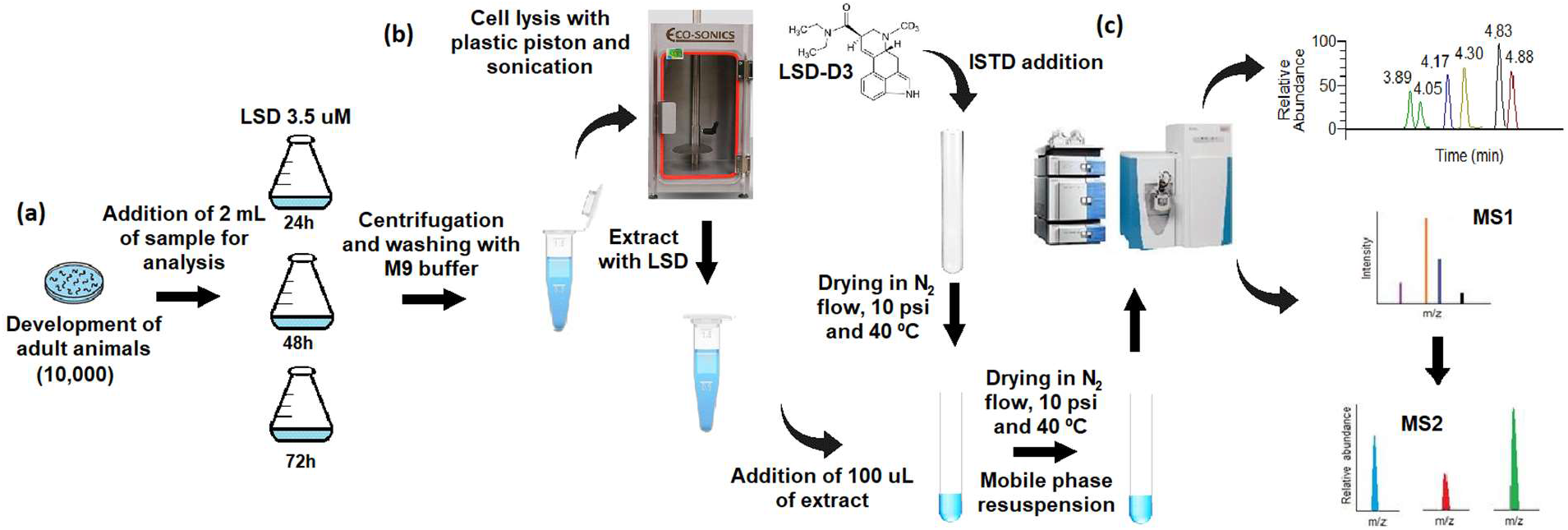
Flowchart with the methodology used in sample preparation.

To obtain homogeneous plates, with the worms at the same stage of development, a synchronization was performed using the bleaching solution technique (1.65% (v/v) NaClO, 0.5 M NaOH). Thus, plates with young adult hermaphrodite worms (in egg laying stage) and eggs were collected with Milli-Q water, and washed 3 times with M9 buffer (20mM KH_2_PO_4_, 0.04M Na_2_HPO_4_, 0.08M NaCl and 0.001M MgSO_4_) to remove any bacterial residue. After the washes, the mixed population of worms and eggs was centrifuged and the bleaching solution added, and incubated for 5 minutes under agitation, for the lysis of the adult worms, however, with the preservation of the eggs. The eggs then were washed to remove bleaching solution and transferred to a new plate without food, for their hatching and development until the first larval stage. Thus, at this stage, the larvae were transferred to a dish with food (bacteria) for the development of the adult stage. In order to obtain a greater number of animals for the preparation of the extract, the eggs of this generation of adults were cultured to obtain a second generation, and these new adults were used for the analyzes (figure 1 (a)).

Before the treatment, synchronized adult animals were collected in liquid media (M9 buffer) and washed 3x to remove bacteria. Each analyzed sample had 10,000 animals. Then, animals were incubated in liquid M9 buffer with LSD at a concentration of 3.5 μM for different timepoints: 24h, 48h and 72h of exposure at room temperature (figure 1 (b)). In addition to the positive points, where LSD was added, for each timepoint, samples containing *C. elegans* incubated in Erlenmeyer flasks without LSD, for the control condition.

After the incubations, control and LSD-treated samples were washed 3x with M9 buffer to remove LSD that wasn’t absorbed. Samples were centrifuged again, supernatant was removed and the pellet was stored at -20°C. To lyse the animals and release the LSD content, sample were thaw, 200 μL of MilliQ water was added to each sample and the animals were macerated with a disposable plastic piston for Eppendorf tubes, using a homogenizer motor, three times for 30 seconds. The sample was then sonicated (Eco Sonics QR350W) for 2 minutes, with pulsator 2 and power 30, on ice. The samples were then centrifuged, the unlysed pellet was discarded, and the supernatant was frozen again at -20°C.

In test tubes, LSD-D3 was added at a concentration of 100 ng/mL and dried in the evaporator with a nitrogen flow at 10 psi and 40º C. Afterwards, 100 μL of the samples, collected in Erlenmeyer flasks after the exposure times, were added to the ISTD-containing tubes and vortexed for 10 seconds. Samples were dried under the same conditions and resuspended in 100 μL of mobile phase 70:30 (water with 0.1% formic acid and 5 mM ammonium formate:methanol with 0.1% formic acid). Finally, the extracts were vortexed for 10 seconds, transferred to vials, and analyzed by UHPLC-MS/MS (figure 1 (c)).

### UHPLC-MS/MS analysis

A Dionex Ultimate 3000 UHPLC system (Thermo Fischer Scientific, Bremen, Germany) coupled to a Q-Exactive hybrid quadrupole-Orbitrap mass spectrometer (Thermo Fischer Scientific, Bremen, Germany) with an ESI source was employed. Chromatographic separation was performed using a Syncronis C18 reversed-phase column (Thermo Fischer Scientific, USA; 1.7 μm, 50 mm x 2.1 mm), with an injection volume of 5 μL at a constant flow rate of 400 μL/min. Mobile phases A (water with 5 mM of ammonium formate) and B (methanol), both containing 0.1% formic acid, were pumped to the mass spectrometer operating in positive ESI mode. The multi-step gradient started with 5% mobile phase B, increasing to 10% at 0.5 minutes, then to 25% at 1 minute and 90% at 6 minutes. After reaching 100% at 8 minutes and maintaining until 9 minutes, the initial chromatographic condition was restored from 9.1 to 11.1 minutes. The ESI source was calibrated daily with a manufacturer’s calibration solution (Thermo Fisher Scientific, Bremen, Germany). Capillary and auxiliary gas temperatures of 380 °C, spray voltage of 3.90 kV and S-lens voltage of 80 V were employed. The nitrogen sheath, auxiliary and sweep gas flow rates were set at 60, 20 and 0 arbitrary units, respectively. Full-scan data were acquired in the range *m/z* 100-1000, at a resolution of 70,000 full width at half maximum (FWHM) with Automatic Gain Control (AGC) of 1×10^6^ and maximum Injection Time (IT) of 100 ms. Parallel Reaction Monitoring (PRM) was performed at a resolution of 70,000 FWHM, AGC of 5×10^5^, maximum IT of 100 ms, loop count of 1 and an isolation window of 1.0 *m/z*. The putative metabolites were selected by an inclusion list for mass fragmentation on a Higher-energy Collisional Dissociation (HCD) with normalized collisional energy (N)CE of 40%. Data were acquired with Thermo Scientific™ TraceFinder™ 4.1 software (Thermo Fisher Scientific, Austin, TX, USA) and analyzed with Xcalibur Qual Browser 4.1 software (Thermo Fisher Scientific, Austin, TX, USA) with ± 5 ppm mass accuracy tolerance.

## Results

To characterize the absorption of LSD by *C. elegans*, an analysis of the extracted ion chromatograms and the MS2 spectra of the LSD absorbed from the prepared extracts and the added ISTD was performed (figure 2). It is possible to evaluate the effectiveness of the ISTD with the non-degradation in the preparation and processing of the samples, as well as the spectrum for identification and characterization of both, LSD and ISTD.

**Figure 2.**
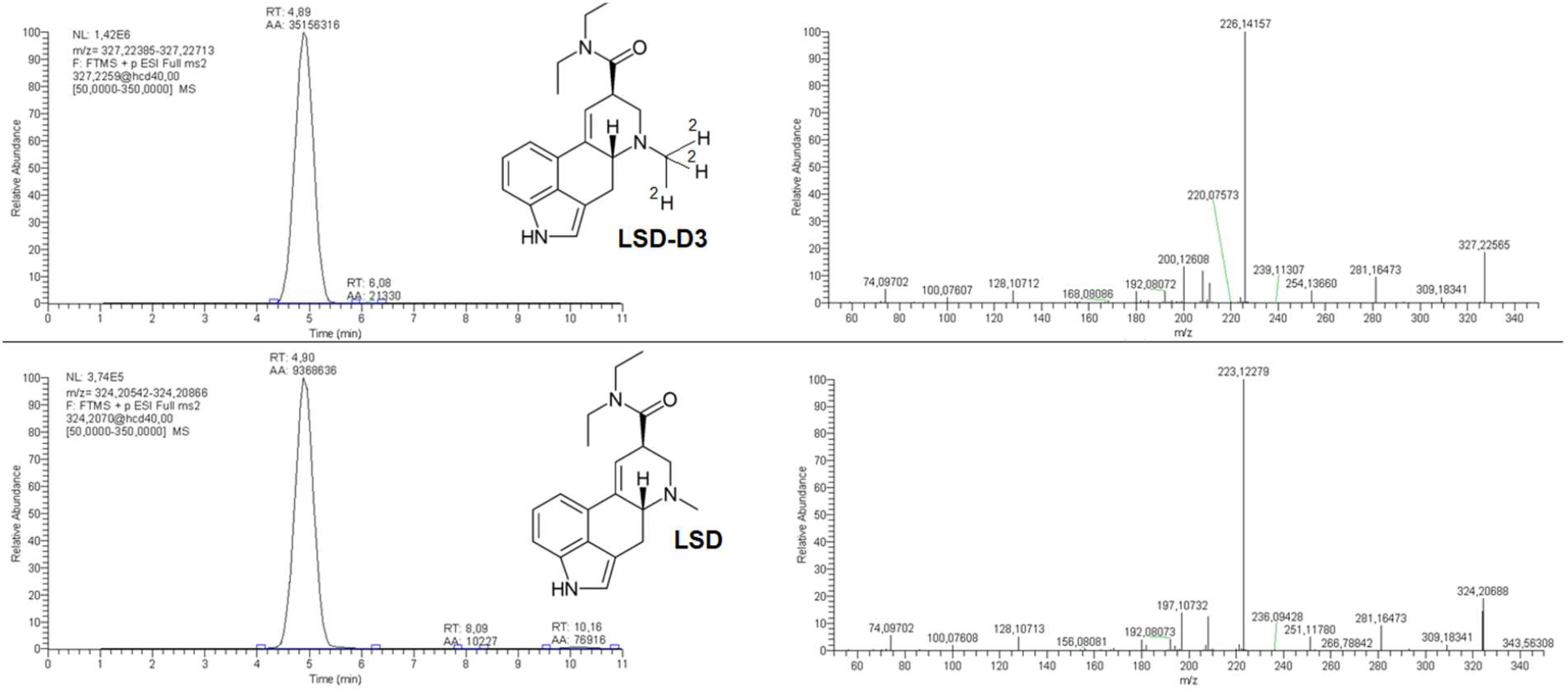
Extracted ion chromatograms and MS2 spectra for ISTD LSD-D3 and LSD from a 24h sample. Analysis of retention times and characteristic fragment ions, with the formation of identity spectra for LSD and for ISTD LSD-D3.

The LSD metabolites identified in the samples by the PRM method at the analyzed degradation times were *nor*-LSD, 2-oxo-3-hydroxy-LSD (O-H-LSD), Lysergic-acid-monoethylamide (LAE) and 13/14-hydroxy-LSD (OH-LSD) (figure 3).

**Figure 3.**
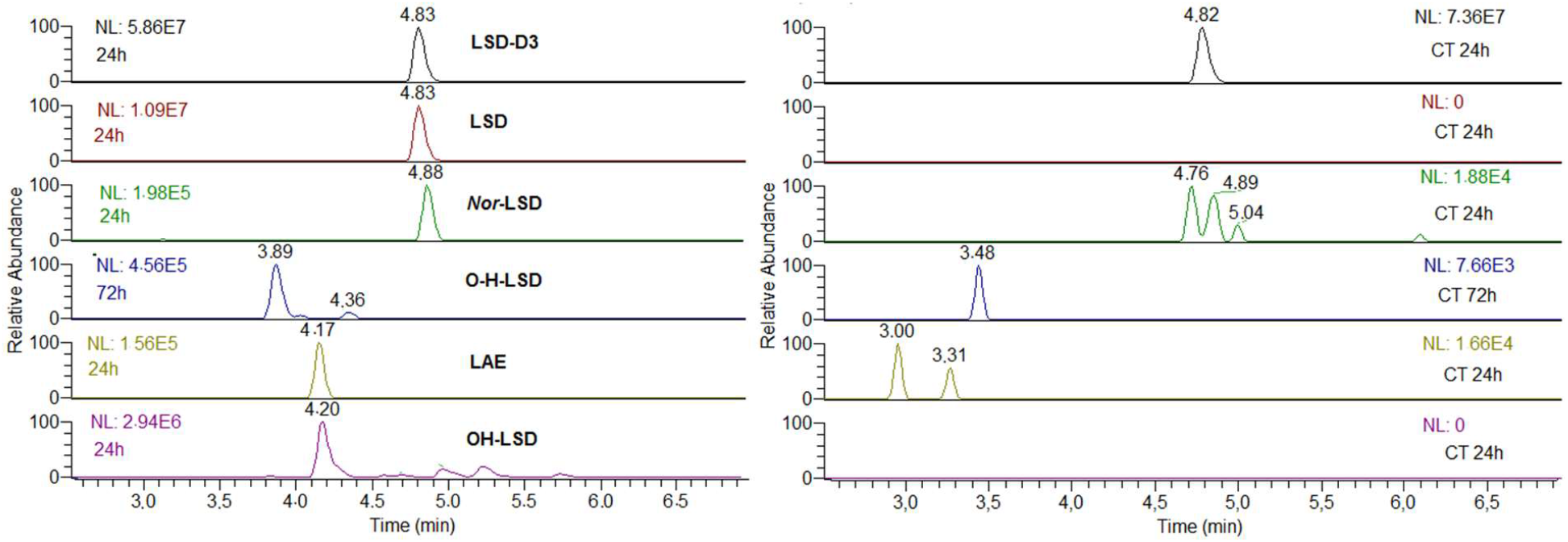
LSD metabolites found in the biological matrix of <u>C. elegans</u>. The chromatograms on the left show the retention times of each metabolite found and the chromatograms on the right the control samples. LSD-D3, LSD, Nor LSD, LAE and OH-LSD in the 24h sample and O-H-LSD in the 72h sample.

The analysis of the MS2 spectra of the metabolites confirm their identities based on the results already published [4,29,33,34], as shown in figure 4. The fragmentation profile of the metabolites is also represented table 1.

**Figure 4.**
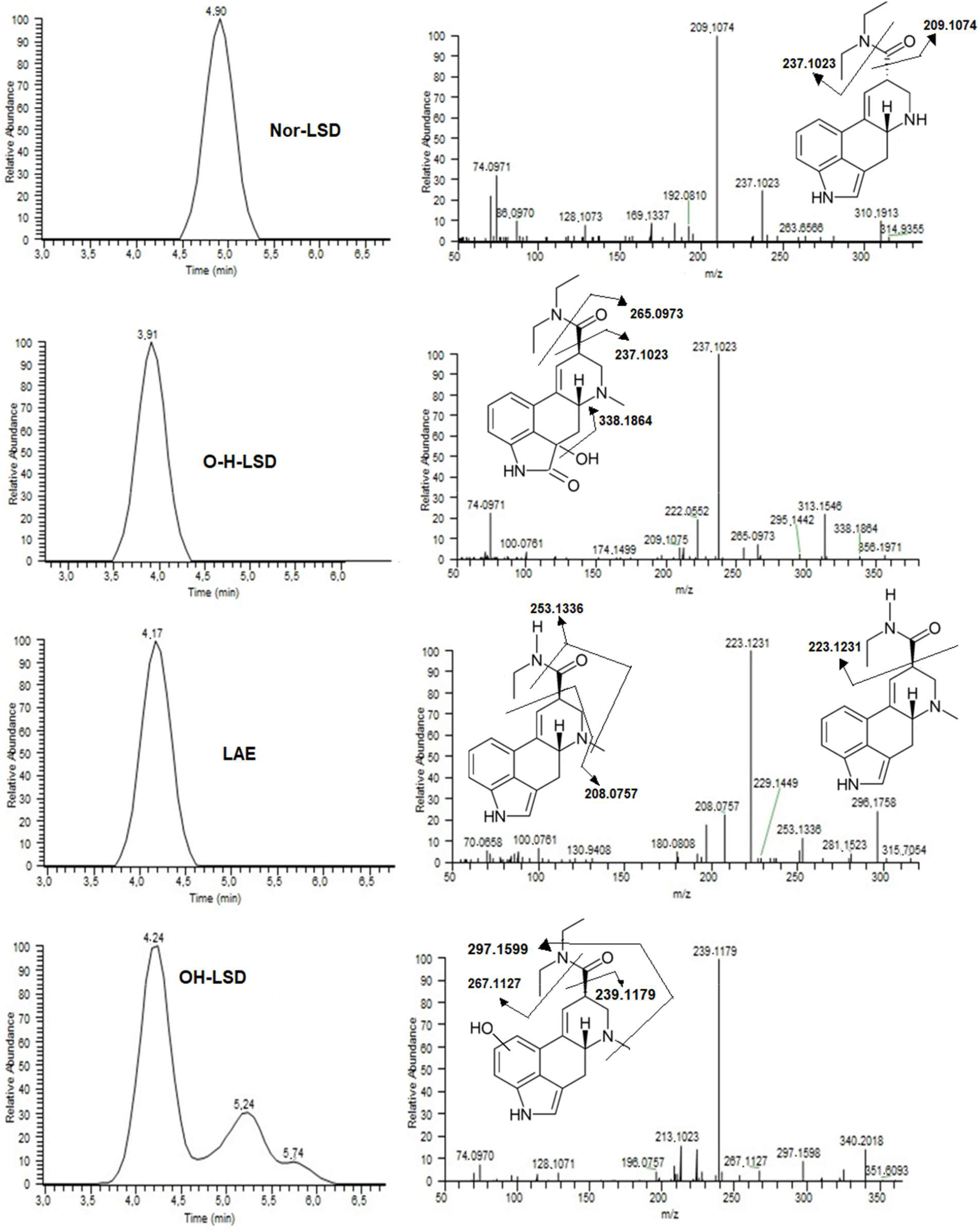
XIC, MS/MS spectra obtained from metabolites and structure with fragmentation patterns.

**Table 1.**
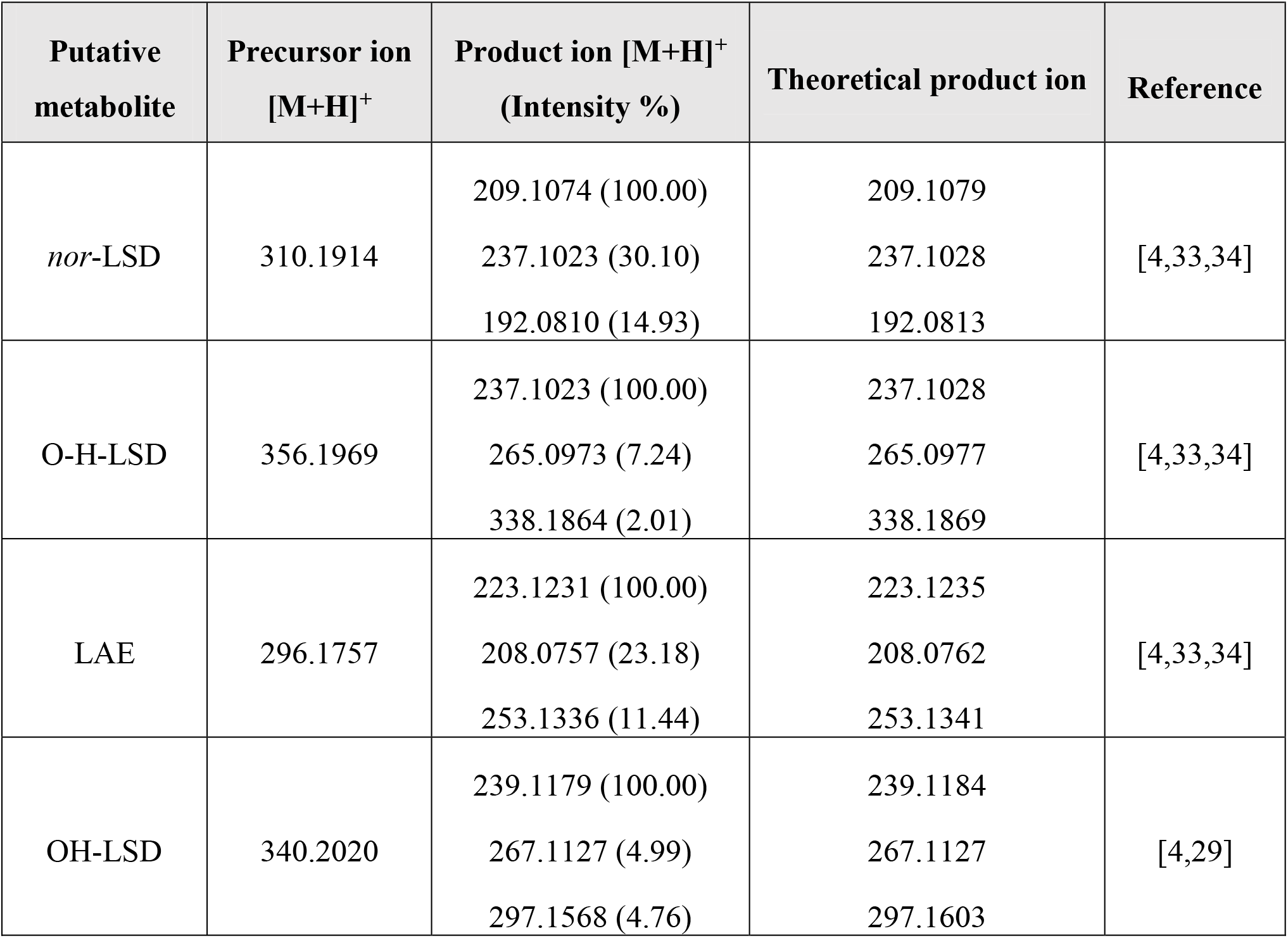
Identity confirmation of the LSD putative metabolites by their fragmentation profile.

With the five identified metabolites, an analysis of their production over time was performed. The obtained metabolic rate is represented in figure 5.

**Figure 5.**
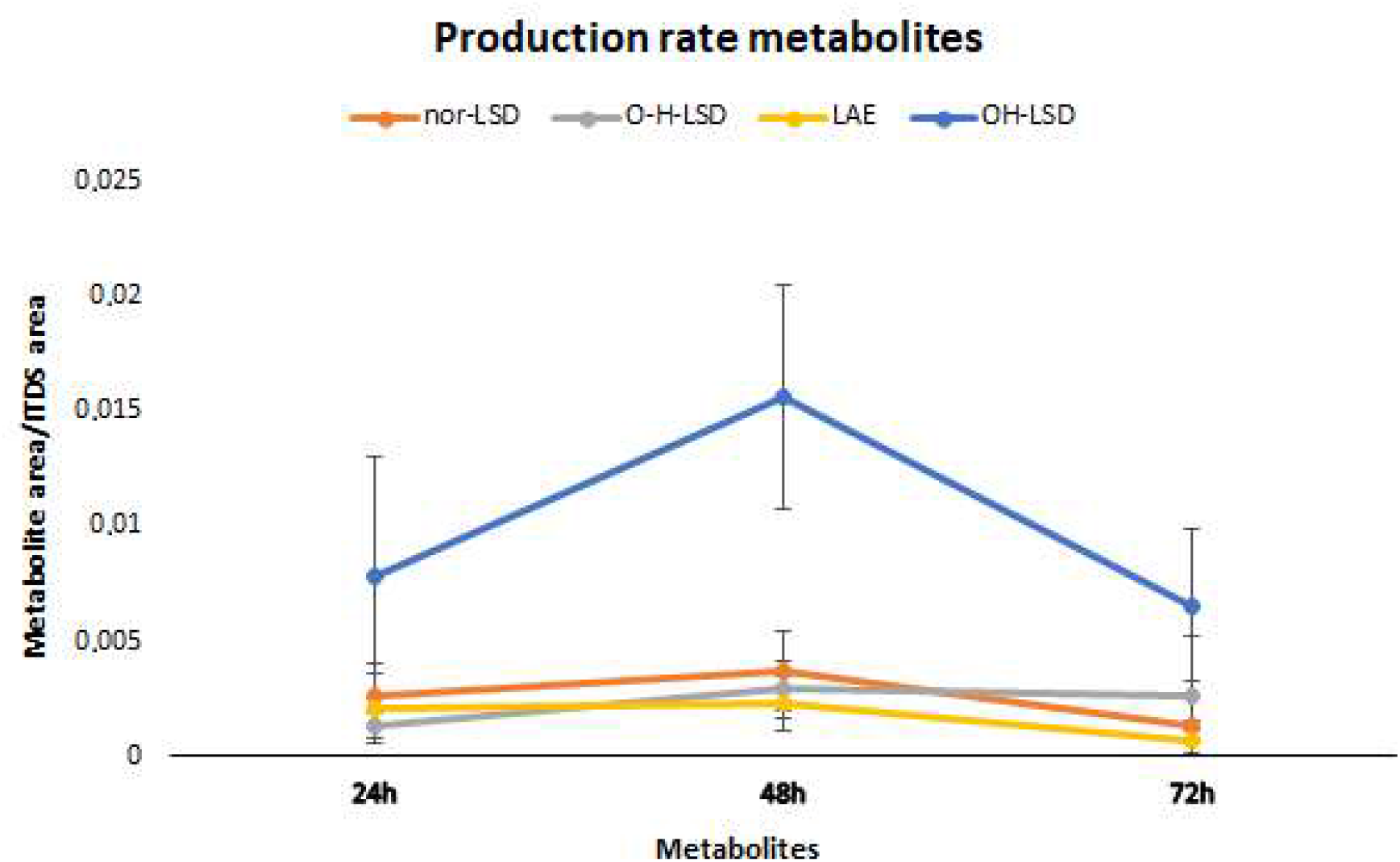
Production rate of LSD metabolites in <u>C. elegans</u>. Analysis of metabolite formation in samples with triplicates at 24h, 48h and 72h. Mean values at each time with the standard deviation.

## Discussion

The rate of metabolic production of LSD varies from species to species, as not all known metabolites were found in the studied species. Thus, there is still no completely defined metabolic pathway [18]. Therefore, the identification of the metabolites is an important step towards the development of a first study on the metabolic effects of LSD in *C. elegans*, as well as the effects on its nervous system. Thus, the PRM method was applied to analyze the MS2 spectra of each metabolite found, in order to form an identification profile, with at least two characteristic fragments for each of them.

*Nor*-LSD is formed by an *N*-demethylation process, with a fragmentation profile showing the transitions *m/z* 310.1914 → *m/z* 209.1074 with loss of diethylamide and *m/z* 310.1914 → *m/z* 237.1023 with loss of diethylamine, both preserving the characteristic *N*-demethylation part. The metabolite O-H-LSD, the main human metabolite found in urine and blood/plasma [29,30], is produced from a hydroxylation and oxidation of the indole moiety of the LSD molecule. Characteristic fragments of *m/z* 237.1023 produced by the loss of diethylamide and water, *m/z* 338.1864 formed by loss of water and *m/z* 265.0973 with loss of diethylamine and water can be observed. LSD also undergoes an *N*-dealkylation reaction to form the metabolite LAE, which presents the fragment *m/z* 223.1231 formed by the loss of ethylamide, *m/z* 253.1336 characterized by the loss of the group (-CH_2_=N-CH_3_), and *m/z* 208.0757 with loss of group (ethylamide-CH_3_). The *m/z* 340.2020 fragment presents the isomers OH-LSD, 2-oxo-LSD and lysergic acid ethyl-2-hydroxyethylamide (LEO), which are formed through hydroxylation of the indole part, oxidation and hydroxylation of diethylamide, respectively. The spectrum with three retention times represents this situation. Due to the hydroxylation suffered by LEO, it can be differentiated from the other two by the spectrum, by the presence of the fragment ion *m/z* 223.1235, with the loss of hydroxylated diethylamide. However, 2-oxo-LSD and OH-LSD cannot be precisely differentiated just by analyzing the mass spectra [4,29]. With the analysis of the structures of the fragments *m/z* 239.1180 (loss of diethylamide), *m/z* 267.1127 (loss of diethylamine) and *m/z* 297.1568 (demethylation plus deethylation), it can be inferred that the highest abundance peak in the sample indicates a hydroxylation on the indole portion of the LSD molecule, possibly corresponding to the metabolite OH-LSD.

Therefore, metabolic alterations in *C. elegans* come from reactions of N-demethylation, *N*-deethylation, hydroxylation and oxidation (figure 6), some modifications also found in metabolic studies performed in humans, rats, mice, cats, and guinea pigs [29,30,33]. Axelrod e*t al*. reported the presence of 2-oxo-LSD metabolized by guinea pig liver microsomes. Slayton and Wright, Szaya and Siddik *et al*. detected the presence of 12-OH-LSD, 13-OH-LSD and 14-OH-LSD in rat bile, respectively, reporting the role of enzymes from the cytochrome P450 family in the formation mechanism [34,35,36,37]. Canezin *et al*. identified and quantified nor-LSD and O-H-LSD in urine samples from two patients, one from a patient who was admitted to the emergency room after ingesting an LSD blotter; the second was obtained from a patient after taking LSD. Furthermore, Canezin *et al*. also identified LEO, LAE and OH-LSD glucuronides in urine samples. Steuer *et al*. detected *nor*-LSD, O-H-LSD, and OH-LSD glucuronide in plasma samples in a study of twelve patients, in addition to LAE and LEO in urine samples. Dolder *et al*. performed two studies with administration of multiple doses of LSD: the first one with sixteen volunteers with a dose of 200 μg, and the second with twenty-four volunteers with a dose of 100 μg. The metabolites O-H-LSD and *nor*-LSD were quantified in urine and plasma samples, as well as the identification of LAE, LEO, 2-oxo-LSD, and OH-LSD glucuronide were performed in a plasma sample. Figure 6 shows the metabolic pathway of LSD in *C. elegans* matrix. In this work, OH-LSD glucuronide was not analyzed, since its main fragment *m/z* 340.2020 is the loss of glucuronide.

**Figura 6.**
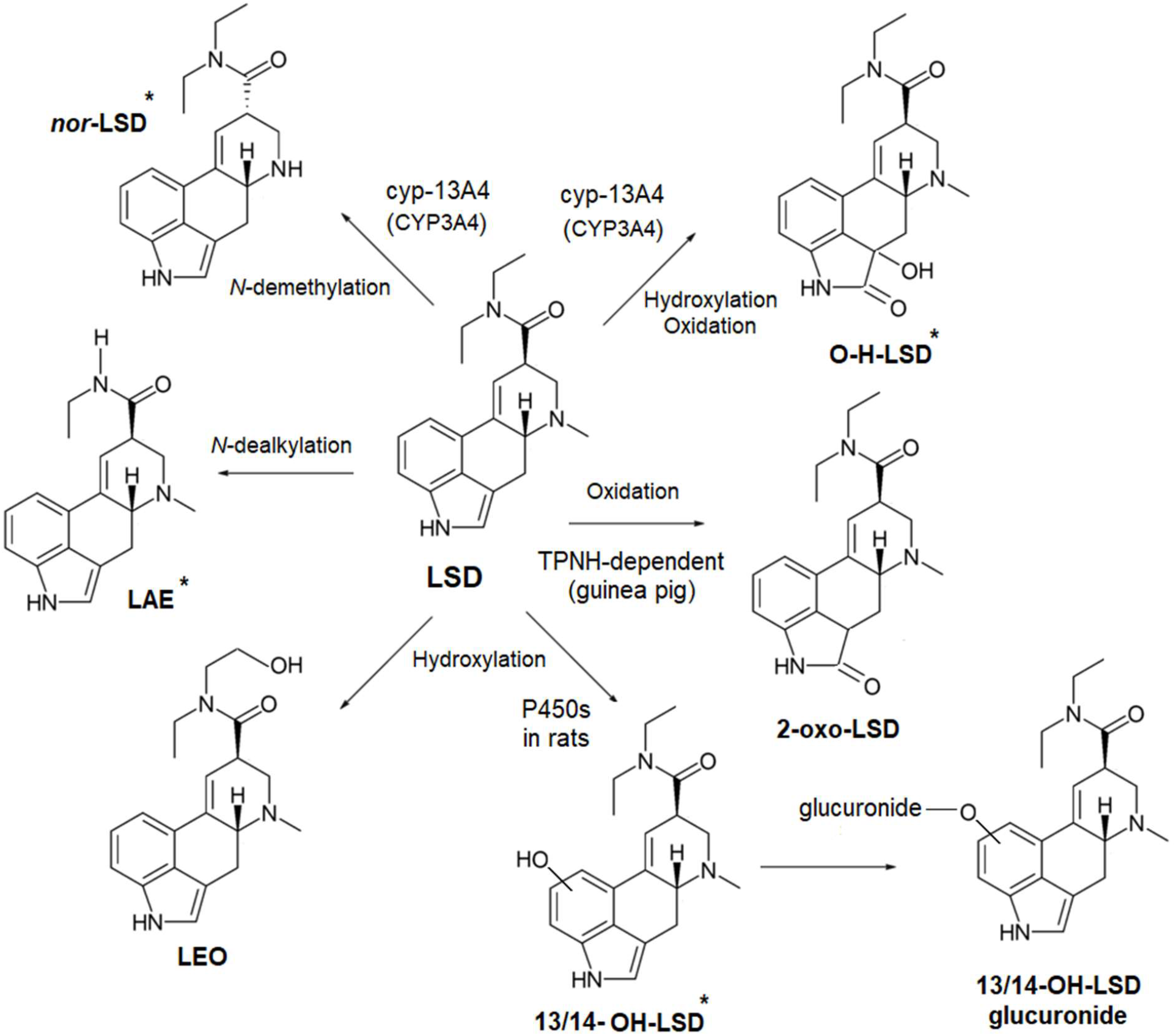
Metabolic pathway of LSD in matrix of <u>C. elegans</u>. All of the above metabolites have been described in humans; 2-oxo-LSD in guinea pigs through the microsomal enzymatic system reduced triphosphopyridine nucleotide (TPNH), 13/14-OH-LSD and 13/14-OH-LSD glucuronide have already been described in rats.* LSD metabolites found for <u>C. elegans</u> in this work. Nor-LSD and O-H-LSD with cyp-13A4 action in metabolism, ortholog of human CYP3A4, one of those responsible for the metabolism in the liver.

Enzyme systems such as cytochrome P450 play an important role in LSD metabolism in humans [29,33]. In this case, CYP1A2, CYP2C9, CYP2E1 and CYP3A4 contribute significantly to the formation of O-H-LSD, while CYP2D6, CYP2E1 and CYP3A4 are significantly involved in the metabolism of LSD to *nor*-LSD [29,33]. According to the genome analysis of *C. elegans* based on data obtained from the Pfam database, cytochrome P450 is listed among the 20 most frequent domains of common proteins [39]. Human cytochromes P450 are divided into 10 clans, while *C. elegans* cytochromes P450 into 4 clans [40]. According to the String homologous protein database, the *C. elegans* cyp-13A4 gene has bilateral orthological level and a similarity (bitscore) of 235.3 with the human CYP3A4 gene [41].

Analysis of the metabolic rate shows a profile of greater intensity of metabolism at 48 hours after absorption and a sharp drop at 72 hours, with the exception of O-H-LSD. Thus, this drop in LSD metabolism after 48 hours can serve as a parameter for future analyzes and studies on metabolic dynamics and kinetics.

## Conclusion

The applied method was effective in detecting LSD metabolites by metabolic degradation in a matrix containing the nematode *C. elegans*, providing for the first time an LSD metabolic pathway for the species. A degradation mechanism was obtained in relation to exposure time, in which five metabolites were found in 24h, 48h and 72h of LSD absorption. The high-resolution analyzes of the spectra allowed identifying the characteristic transitions in a sensitive and efficient way, providing a profile of identification of metabolites through the obtained MS2 spectra. The results pave the way for future studies on the dynamics of the metabolites and their mechanism of action in the matrix of *C. elegans*.

## Acknowledgements

This project was partially supported by the Beckley Foundation and an intramural grant from D’Or Institute for Research and Education. The authors thank FAPERJ (process number SEI-260003/015706/2021 – APQ1), Institute of Biomedical Sciences, Federal University of Rio de Janeiro (UFRJ), Rio de Janeiro, Rio de Janeiro, Brazil, CAPES (phD scholarship: 88887.600288/2021-00; master’s scholarship: 88887.827807/2023-00; master’s scholarship: 88887.713023/2022-00) and CNPq (phD scholarship: 162996/2018-7).

