## Supplementary material for "Systematic Characterization of LSD metabolites in *C. elegans* by ultra-high performance liquid chromatography coupled with high-resolution tandem mass spectrometry": Table 1: Table 1.docx

**Table 1.** Identity confirmation of the LSD putative metabolites by their fragmentation profile.

| **Putative metabolite** | **Precursor ion [M+H]^+^** | **Product ion [M+H]^+^ (Intensity %)** | **Theoretical product ion** | **Reference** |
| --- | --- | --- | --- | --- |
| *nor*-LSD | 310.1914 | 209.1074 (100.00)  237.1023 (30.10)  192.0810 (14.93) | 209.1079  237.1028  192.0813 | [4,33,34] |
| O-H-LSD | 356.1969 | 237.1023 (100.00)  265.0973 (7.24)  338.1864 (2.01) | 237.1028  265.0977  338.1869 | [4,33,34] |
| LAE | 296.1757 | 223.1231 (100.00)  208.0757 (23.18)  253.1336 (11.44) | 223.1235  208.0762  253.1341 | [4,33,34] |
| OH-LSD | 340.2020 | 239.1179 (100.00)  267.1127 (4.99)  297.1568 (4.76) | 239.1184  267.1127  297.1603 | [4,29] |
